# Convergent evolution of mevalonate pathway in *Inonotus obliquus* and *Betula pendula*

**DOI:** 10.1101/2021.11.28.470225

**Authors:** Omid Safronov, Guleycan Lutfullahoglu Bal, Nina Sipari, Maya Wilkens, Pezhman Safdari, Olli-Pekka Smolander, Jenna Lihavainen, Niko Silvan, Sitaram Rajaraman, Pia K. Laine, Lars G Paulin, Petri Auvinen, Tytti Sarjala, Kirk Overmyer, Jaakko Kangasjärvi, Brendan Battersby, Uwe Richter, Jarkko Salojärvi

## Abstract

*Inonotus obliquus*, Chaga mushroom, is a fungal species from *Hymenochaetaceae* family (*Basidiomycota*) which has been widely used for traditional medicine in Europe and Asia. Here, chaga genome was sequenced using Pacbio sequencing into a 50.7Mbp assembly consisting of 301 primary contigs with an N50 value of 375 kbp. Genome evolution analyses revealed a lineage-specific whole genome duplication event and an expansion of Cytochrome P450 superfamily. Fungal biosynthetic clusters were enriched for tandemly duplicated genes, suggesting that biosynthetic pathway evolution has proceeded through small-scale duplications. Metabolomic fingerprinting confirmed a highly complex terpene biosynthesis chemistry when compared against related fungal species lacking the genome duplication event.

## Introduction

*Inonotus obliquus*, Chaga mushroom, is a fungal species from *Hymenochaetaceae* family (*Basidiomycota*) distributed across the boreal forest zone in the Northern hemisphere. It causes aggressive white rot disease mainly among *Betula* family members (Blanchette, 1982), but upon suitable conditions it can infect also other tree species such as oaks, poplars, ashes and maples (Ryvarden & Gilbertson, 1993). White rot disease is the result of lignin degradation (having darker color) while the light-coloured cellulose is left intact. The infection starts when *I. obliquus* spores get access to the hardwood of the stem through an opening or wounded bark. At the later stages of the infection, *I. obliquus* appears as a sterile conk, a solid charcoal-black mass on the surface of bark (Blanchette, 1982). The sterile conk has been used in traditional medicine in many cultures. A large body of research on biochemical compounds extracted from the conk suggests that the species may have a wide range of pharmaceutical, medicinal, and industrial applications (Ma, Chen, Dong, & Lu, 2013; Nagajyothi, Sreekanth, Lee, & Lee, 2014; Song, Liu, Kong, Chang, & Song, 2013; Yan et al., 2014).

Betulin (BE) and betulinic acid are highly abundant triterpenoids in the bark of all birch family members involved in protection against fungi, bacteria and viruses; they collectively form 30-60% of total tissue composition, depending on the species and the tissue type (Holonec, Ranga, Crainic, Truţa, & Socaciu, 2012; P. Kovalenko et al., 2009; Safronov et al., 2019). Both betulinate compounds are being studied for industrial applications (Šiman et al., 2016), and as therapeutic substances in oncology (Król, Kiełbus, Rivero-Müller, & Stepulak, 2015) and infectious diseases (fungal, bacterial, and viral infections) (Gong et al., 2004; Salin et al., 2010; Shai, McGaw, Aderogba, Mdee, & Eloff, 2008). In plants, the biosynthesis of betulinate compounds starts with squalene, a product of mevalonate pathway, and involves two enzymatic steps where squalene is first converted to lupeol via lupeol synthase and then to betulinate by lupeol monooxygenase, an enzyme which is a member of the large family of cytochrome P450 monooxygenases, more specifically subfamily 716 (CYP716). Betulin biosynthesis is found across a wide taxonomic range in plants, from *Malvales* (H. J. Zhang et al., 2003), *Fagales* (Safronov et al., 2019), *Rosales* (Andre et al., 2013; S. Zhao et al., 2015), *Fabales* (Wu, Niu, Bakur, Li, & Chen, 2017), *Vitales* (Fukushima et al., 2011), and *Asterales* (Siddiqui et al., 2019) to *Arecales* (Khelil, Jardé, Cabello-Hurtado, Ould-el-Hadj Khelil, & Esnault, 2016; Koolen et al., 2012), suggesting either ancestral origin or convergent evolution. A comparative genomic analysis of the bark tissue in silver birch (*Betula pendula*) and grey alder (*Alnus glutinosa*) revealed birch-specific evolution of mevalonate pathway (MVA), where a tandem duplication of lupeol synthase colocalized with lupeol 28-monooxygenase was suggested as the reason for increased production of betulinate compounds in birch phellem (Safronov et al., 2019). Interestingly, in addition to plants, betulinate compounds have also been identified in diverse range of fungal species from *Eurotiales* (Khouloud Barakat, 2016), *Hymenochaetales* (Yin, Cui, & Ding, 2008), and *Polyporales* (Alresly et al., 2015) families, even though no members of CYP716 gene family have yet been identified or characterized in fungi. Birch fungal pathogens *Inonotus obliquus* and *Fomitopsis betulina* are such examples of fungal species that produce BE and BA compounds (Alresly et al., 2015; Yin et al., 2008), even though the compounds are anti-fungal by nature. The evolution of betulinate biosynthesis in these two fungal species is not known, but one can hypothesize it to be the result of either convergent evolution or horizontal gene transfer (HGT) of the responsible cytochrome P450 monooxygenase enzymes from the host species. There exists a recent study on the diversification and distribution of CYP716 enzyme in eudicots (Miettinen et al., 2017), but no studies of this enzyme in fungal species have been carried out, and the enzymes underlying betulin production in the fungal species, known to produce betulinate compounds, have not yet been identified.

The CYP450 monooxygenase enzymes are among the oldest and largest gene families, encompassing both prokaryotic and eukaryotic organisms (Sezutsu, Le Goff, & Feyereisen, 2013). They act as key enzymes for detoxification of toxic compounds, and they have an important function in secondary metabolism related to adaptation to environmental conditions. The low sequence similarity, high functional diversity and enzymatic promiscuity among CYP450 monooxygenase enzymes makes functional predictions difficult. The CYP450s are generally classified into families and subfamilies based on sequence similarity; the sequences with identity >40% are assigned into families and sequences with >55% similarity into their own subfamilies; novel candidates with lower identity to the set of identified CYP450s form new candidate families. Based on these criteria, so far over 800 different CYP families have been identified (Lepesheva et al., 2008).

In this study we sequenced and assembled an *I. obliquus* genome from an isolate from Merikarvia region in Finland. The genome was annotated using *ab initio* gene model prediction and spliced transcript data obtained from total RNA sequencing. We carried out comparative genomic and gene family expansion analysis among 16 *Basidiomycete* and 3 *Ascomycetes* species together with *I. obliquus* genome and studied the untargeted terpenoid metabolic fingerprints (using UPLC-QTOF/MS) in five strains of *I. obliquus* and one *Fomitiporia mediterranea* strain, focusing on the quantification of betulin (BE) and betulinic acid (BA) abundances across the samples. To confirm our functional predictions we cloned the candidate lupeol synthase from *B. pendula* and CYP450 monooxygenase enzymes from both *B. pendula* and *I. obliquus* and tested their ability to produce betulin compounds.

## Materials and methods

### Sample collection

Four *Inonotus obliquus* strains (Supp. table 1) were collected and isolated from different regions in Finland and one from Altai mountains in Russia. The strain from Merikarvia was selected for whole genome sequencing (location 61°58’38.6”N 21°44’43.1”E). In addition, we also obtained a strain of *Fomitiporia mediterranea* as an outgroup to chaga (Mycobank: MB384943). All samples were cultivated on Hagem agar overlayed by a cellophane membrane.

The isolation of chaga mushrooms from the host trees was done by cutting a piece of the conk (Supp. Fig 1), which was then laid on agar plate after short H_2_O_2_ bath. The samples were re-cultured repeatedly and sequenced for internal transcribed spacer 1 (ITS1) [TCCGTAGGTGAACCTGCGG] and ITS4 [TCCTCCGCTTATTGATATGC] regions confirm the species assignment of *I. obliquus* isolate.

### RNA isolation, sequencing, and *de novo* assembly of transcriptome

To isolate the total RNA from *I. obliquus,* the method from Chang *et al.* (Chang, Puryear, & Cairney, 1993) was used. Briefly, the *I. obliquus* was inoculated and grown on autoclaved wood dust from a clone of *B. pendula* (12 years old tree, 167 cm^2^ disk, dry weight of 200 grams) sequenced for *B. pendula* reference genome (Salojärvi et al., 2017). A total of 150 milligrams of ground sample (mortar and pestle, and liquid N_2_) was transferred on ice for 30 seconds, and 500 μl of pre-warmed (+65-68°C) extraction buffer (2% CTAB, 2% PVP K-30, 100 mM Tris-HCl [pH 8.0], 25 mM EDTA, 2 M NaCl, and 200 μl β-MeOH/10 ml of extraction buffer) was added and vortexed vigorously. Extraction was carried out three times with chloroform:isoamyl alcohol (24:1) by spinning at 200-300 rpm for 15 minutes, and then centrifuging at 10 000 rpm 15 minutes. Then, 1/4 volume 10 M LiCl was added and left to precipitate on ice overnight. The overnight sample was centrifuged with 10000 rpm for 20-30 minutes at +4°C, and the resulting pellet was dissolved in 500 μl of pre-warmed (+65°C) sodium dodecyl sulfate–Tris-HCl–EDTA (SSTE) buffer, and extracted once (or several times, if necessary) with chloroform:isoamyl alcohol (24:1). The mixture was precipitated by adding 2 volumes of absolute EtOH (place at −20°C overnight), and centrifuged at 13 000 rpm, for 20-30 minutes at +4°C. the precipitate was washed with 70% EtOH, after which the pellet was dried, and then dissolved in 10-30 μl RNase-free water, and RNase inhibitor was added.

TruSeq stranded mRNA kit was used to construct the RNA-seq library. The cDNA was synthesized from 5 μl of total RNA extracted from reference *I. obliquus* plate using random hexamers. DNA polymerase I and dUTP nucleotides were used to synthesize the second strand of cDNA. Then, double stranded cDNA were purified, and ends were repaired. Library preparation was continued by A-tailing, and ligation of Y-adaptors containing indexes from the kit. The fragments were amplified using polymerase chain reaction (PCR), followed by purification steps using AMPure XP. The sequencing was carried out in HiScan SQ platform (paired-end 88 bp + 74 bp).

The raw paired end RNA-seq data were controlled for quality using FastQC v0.11.2 (Andrews). Trimmomatic v0.33 (Bolger, Lohse, & Usadel, 2014) was used in pair-end mode to remove the adapters, barcodes, low quality bases from both ends of each sequence, and reads shorter than 25 base pairs (LEADING:20, TRAILING:20, MINLEN:25, -phred33). After the removal of duplicate sequences, the unpaired sequences were mapped to *I. obliquus* reference genome using Tophat2 (Kim et al., 2013) for junction discoveries (-i:10, and --coverage-search); paired end reads were mapped separately (Tophat2; -i:10, and --coverage-search). The aligned reads were separated according to their orientation on reference genome to forward and reverse strands, which were then aligned individually by Trinity v2.1.1, using --genome_guided_bam, and --genome_guided_max_intron: 1 000 options (Grabherr et al., 2011) for *de novo* transcriptome assemblies. The forward and reverse *de novo* transcriptome assemblies were combined, and duplicated assemblies were removed using GenomeTools v1.5.1, using sequniq option (Gremme, Steinbiss, & Kurtz, 2013). The unique *de novo* transcriptome assemblies were clustered by using CD-HIT v4.6 (Godzik & Li, 2006) and aligned to *I. obliquus* reference genome by Program to Assemble Spliced Alignments (PASA v2.2.0) (Brian J. Haas et al., 2008).

The processing of the publicly available RNAseq data (Fradj et al., 2019) was carried out in a similar manner. Both data sets were mapped to *I. obliquus* gene models using kallisto quant v0.44.0 (Bray, Pimentel, Melsted, & Pachter, 2016). The orphan reads and pair-end reads (separated during preprocessing by trimmomatic) were mapped separately by using kallisto quant single (options: -- single, -l 200, -s 20, -b 4000) and pair-end (option: -b 4000) modes, respectively. The raw count table from Kallisto was imported to R (for both single and pair-end count tables) using tximport package v1.18.0 with default options (Soneson, Love, & Robinson, 2015). The single and pair-end counts were summed together to form a single count table for each data set. Differential gene expression analysis was conducted using DESeq2 (Love, Huber, & Anders, 2014). The final tables for differentially expressed genes (DEg) were filtered based on the false discovery rate adjusted p-value threshold of 0.05 (p-adj. ≤ 0.05).

### DNA isolation, genome assembly and annotation

Modified version of Lodhi *et al.* (Lodhi, Ye, Weeden, & I. Reisch, 1994) was used for DNA extraction from *I. obliquus* strains. Maximum 0.5 g of material was ground in liquid N_2_. The ground sample was transferred into ice cold Sodium chloride-Tris-EDTA (STE) buffer (1,4 M NaCl, 0 mM EDTA, 100 mM Tris-HCl pH 8.0), and centrifuged for 5 minutes at 8 000 rpm and +4°C. STE buffer was discarded, and 10 ml of pre-warmed (60°C) cetyltrimethyl ammonium bromide (CTAB) buffer (1 liter CTAB: 20 mM EDTA, 100 mM Tris-HCl pH 8.0, 1.4 M NaCl, 2.0% CTAB, 1.0% PVP 40, and 2% β-MeOH [50μl]) was added to pellet. Subsequently, the mixture was vortexed and incubated for 30-60 min at 60°C and cooled to the room temperature. Chloroform: isoamyl alcohol (IAA) (24:1 ration) mixture was added for extraction (centrifugation: 15 minutes, 10000 rpm at room temperature). The supernatant was collected to a new tube and mixed with 2X CTAB buffer, which was then vortexed and incubated for 30-60 min at 60°C. The chloroform:IAA extraction step was repeated 2-3 times, followed by adding of 2X volume of cold (−20°C) absolute ethanol (EtOH) to supernatant. The EtOH mixture was stored for overnight at +4°C. The mixture then was centrifuged for 15 minutes, at 10000 rpm and 4°C. DNA pellet was washed with absolute EtOH (−20°C) and air dried. The sample was treated for the RNA (RNase A), followed by chloroform:IAA extraction, EtOH precipitation, air drying of the DNA pellet, dissolving in DNase/RNase free water, and storing at −80°C.

The genome of the *Inonotus obliquus* was sequenced with Pacific Biosciences PacBio RSII instrument using P6-C4 chemistry. Eight SMRTcells were used for sequencing the sample with movie time of 240 minutes. The number of obtained sequences was 712,759 which totaled up to 4.82 Gb of data with read length N50 of 9200 bp. At first, hierarchical Genome Assembly Process (HGAP) V3 implemented in SMRT Analysis package (v2.3.0) was used to generate an initial *de novo* genome assembly with default parameters. Mitochondrial genome contig was separated from the chromosomal contigs and circularized manually using GAP4 program (Bonfield, Smith, & Staden, 1995). Obtained mitochondrial sequence in length of 118 085 bp and > 4000X sequencing coverage was polished using SMRT Analysis RS Resequencing protocol with Quiver consensus algorithm. Second, the FALCON assembly program (Chin et al., 2016) was used to generate the final *de novo* genome assembly with seed read length of 10 000 bp. Obtained contig sequences were polished using SMRT Analysis RS Resequencing protocol with Quiver consensus algorithm with approximately 75x coverage. To quantify completeness of the genome, BUSCO (v3.0, Fungi datasets, -m geno, – long) (Waterhouse et al., 2017) was used.

Repeat analysis of the contigs was carried out according to the guidelines of RepeatModeler and RepeatMasker (http://www.repeatmasker.org/, v 4.0.7). To predict the gene models, multiple evidence tracks from different platforms were obtained: *ab initio* gene predictors based on Hidden Markov Models (HMMs), spliced transcript evidence from RNA-seq, and orthologous proteins from closely related fungal species. HMM-based models such as AUGUSTUS (v3.3.2) (Stanke & Morgenstern, 2005), and GeneMark-ES (version 4.33; --fungus mode, and –evidence: *de novo* transcriptome assembly) (Besemer & Borodovsky, 2005) were used for *ab initio* gene predictions. In addition, BRAKER2 (Hoff, Lange, Lomsadze, Borodovsky, & Stanke, 2015) (options: --fungus, -- rounds=100, and --bam) was run for *ab initio* gene predictions. To identify the open reading frames (ORFs) within the genome, getorf (EMBOSS v6.6.0.0) program (Rice, Longden, & Bleasby, 2000) was used (-find:1, and –maxsize: 5000). The ORFs were then queried against NR database by DIAMOND (v0.9.24, blastp, --more-sensitive) (Buchfink, Xie, & Huson, 2014) and filtered for similarity (sequence identity ≥ 75, and score ≥ 300); the homologous sequences above the threshold were collected. Selected ORFs were used as the input for exonerate (v2.46.2, --model:protein2genome, --minintron:10, --maxintron:1000; --percent:65) (Slater & Birney, 2005) to map the candidate ORFs to *I. obliquus* reference genome. Additionally, orthologous proteins from 13 fungal species (*Coprinopsis cinerea*, *Fomitiporia mediterranea*, *Heterobasidion annosum*, *Laccaria bicolor*, *Onnia scaura*, *Phanerochaete chrysosporium*, *Phellinus ferrugineofuscus*, *Porodaedalea niemelaei*, *Postia placenta*, *Puccinia graminis*, *Rickenella mellea*, *Schizopora paradoxa*, *Trichaptum abietinum*) were aligned against *I. obliquus* reference genome with exonerate (v2.46.2, --model:protein2genome, -- minintron:10, --maxintron:1000; --percent:65) (Slater & Birney, 2005). In addition to orthologous proteins, the protein sequences discovered from BUSCO predictions were collected and aligned to reference genome by exonerate as well using the same parameters as given above (Slater & Birney, 2005). All the evidence (*ab initio* gene models, spliced transcript alignments, spliced protein alignments, ORFs, and BUSCO) was combined to consensus, high-confidence gene models, using EVidenceModeler (v1.1.1). This was followed by the addition of untranslated regions (UTR) to the gene models by PASA (Brian J. Haas et al., 2008).

Mitochondrial genome was also assembled and annotated as described previously (Salojärvi et al., 2017), resulting in 29 tRNAs, 32 coding sequences, and 3 rRNAs.

Interproscan (v5.25-64.0) (Quevillon et al., 2005) was used to assign the protein function to gene models. Additionally, Ensemble Enzyme Prediction (E2P2, v3.1) (Schlapfer et al., 2017) and antiSMASH (v2.0) fungal version (Blin et al., 2013) were used to predict the metabolomic pathways.

### Comparative genomic analyses

The proteomes of twenty fungal species *Laccaria bicolor, Coprinopsis cinerea, Schizophyllum commune, Fomitiporia mediterranea, Inonotus obliquus, Onnia scaura, Phellinidium ferrugineofuscum, Porodaedalea niemelaei, Trichaptum abietinum, Rickenella mellea, Schizopora paradoxa, Fomitopsis betulina, Postia placenta, Phanerochaete chrysosporium, Puccinia graminis, Heterobasidion annosum, Ustilago maydis, Saccharomyces cerevisiae, Schizosaccharomyces pombe,* and *Neurospora crassa* from Ascomycete and Basidiomycete clades were downloaded from MycoCosm (https://mycocosm.jgi.doe.gov) and included for gene family analysis by Orthofinder (Emms & Kelly, 2015) (v2.3.3), run with default parameters.

### Synteny analyses

Synteny analysis of self-self alignment of *I. obliquus*, and four other fungal species, namely *F. mediterranea*, *S. paradoxa*, *F. betulina*, and *P. niemelaei,* were conducted using SynMap application in CoGe platform (https://genomevolution.org/coge/), using Quota Align algorithm with default parameters. The list of syntenic duplicates were obtained from DAGchainer (B. J. Haas, Delcher, Wortman, & Salzberg, 2004); tandem duplicates were obtained as part of the preprocessing pipeline.

### Discovery of secreted proteins and carbohydrate active enzymes (CAZymes)

Getorf function of EMBOSS (v6.6.0.0) (Rice et al., 2000) was used to discover the ORFs (-find:1, and –maxsize: 1000). All the ORFs were analyzed by signalp (v5) (Almagro Armenteros et al., 2019) for the presence of signal peptide. Signal peptides were removed from predicted ORF sequences, and the cysteine amino acids were counted for every sequence. ORF sequence with three cysteine residues was predicted as a possible secreted protein (SP).

CAZymes are annotated during gene model annotation steps. In order to further classify these enzymes, total proteome of *I. obliquus* was queried against dbCAN2 database using DIAMOND (v0.9.24, blastp, --more-sensitive) (Buchfink et al., 2014; H. Zhang et al., 2018), and the best hit was selected (score ≥ 200, percentage identity ≥ 55) as a homologous sequence.

### Gene tree of cytochrome P450 monooxygenases

A phylogenetic tree was constructed for 15 CYP716 gene models from Streptophyta species which have been confirmed to produce betulinate compounds [*Betula pendula* (Salojärvi et al., 2017), *Betula platyphylla*, *Phoenix dactylifera* (Al-Mssallem et al., 2013), *Medicago truncatula* (Tang et al., 2014), and *Vitis vinifera* (The French–Italian Public Consortium for Grapevine Genome et al., 2007)], after which CYP450 genes with high sequence similarity (< 40%) in *I. obliquus* and *F. betulina* were added to the tree. A total of 77 sequences were subjected to multiple sequence alignment (MSA) using MUSCLE (v3.8.31, -maxiters: 1000) (Edgar, 2004). The amino acid sequences were reverse translated by PAL2NAL (Suyama, Torrents, & Bork, 2006), and both amino acid and nucleotide sequences were used to construct the phylogenetic trees by RaxmlHPC-HYBRID-AVX2 (v8.2.12) (Stamatakis, 2014).

### Mass spectrometry, sample preparation and derivatization of *I. obliquus* and *F. mediterranea* metabolites

Five strains of *I. obliquus* and one *F. mediterranea* (three biological replicates for each) were grown in liquid Hagem media for two weeks. Submerged myceliums were washed with sterile milli-Q water three times, and grinded with liquid nitrogen. All samples (fresh wight ~500mg, dry weight ~30mg) were extracted twice with 1.0 ml of ethyl acetate (Merck) by vortexing for 15 min and centrifuged for 10 min in 15000 rpm according to Cao et al. (G. Zhao, Yan, & Cao, 2007) at room temperature. Internal standards (ISTD) (10 μl, 10 μg/ml), testosterone and 4-methylumbelliferone (Sigma), were added to each sample in the first extraction step. The supernatant was evaporated to dryness with MiVac Duo concentrator +40°C (GeneVac LtD, Ipswich, UK) and the residue was re-solubilized in 100 μl ACN (Honeywell). Quality control (QC) sample was prepared by combining extracts from each sample line.

The triterpenoid profiling was executed from the extracts with UPLC-PDA-QTOF/MS. The UPLC-MS system consisted of a Waters Acquity UPLC attached to a Acquity PDA-detector and to a Waters Synapt G2 (HDMS) QTOF mass spectrometer (Waters, Milford, MA, USA). The separation of the analytes was executed in Acquity BEH C18 (2.1mm x 50mm, 1.7μm) column (Waters, Milford, MA, USA) with the temperature of +40°C. The autosampler temperature was set to +27°C. The mobile phase consisted of water (A) and acetonitrile (B) both with 0.1% formic acid and the flow rate was 0.6 ml/min. The injection volume was 3 μl. The linear gradient started with 30% B and proceeded to 98% in 9 min, followed by 1 min at 98% B, giving a total run time of 10 min. ESI/MS detection was performed in positive sensitivity ion mode with capillary voltage 3.0 kV, cone voltage 30 V, desolvation gas 800 L/h, cone gas 20 L/h, desolvation temperature 320 °C, source temperature 120 °C and extractor lens 3.00 V. The MarkerLynx software (Waters, Milford, MA, USA) was used for data processing. UV spectra, negative MS-runs and fragmentation patterns from MS^e^ runs of QC sample were used as additional tools for annotation of triterpenes, sterols and phenolic compounds in *Inonotus obliquus* samples.

The standard solutions of betulin and betulinic acid (1.0 mg/ml) were prepared in ethyl acetate and testesterone (100 μg/ml) standard was prepared in methanol. Working solutions (10 μg/ml, 100 ng/ml) were prepared by diluting the standard solution with acetonitrile. The optimization of quantification method was executed with a mix of betulin, betulinic acid and testosterone standards (100 ng/ml). Standard mix (100 μl) was derivatized with PTSI, and MS parameters were optimized by repeated injection of the sample.

Due to very low concentration of betulin and betulinic acid, extracts were derivatized with p-toluenesulfonyl isocyanate (PTSI) (Hu et al., 2013; Zuo, Gao, Liu, Cai, & Duan, 2005) to improve sensitivity. After metabolite profiling with UPLC-QTOF/MS, samples (90 μl) were derivatized for 3 min with 10 μl of 60% p-toluenesulfonyl isocyanate (PTSI) (Sigma) in acetonitrile (Hu et al., 2013; Zuo et al., 2005). The derivatization reaction was terminated with 50 μl of methanol (Merck) with 30 s vortex mixing, giving the total volume of 150 μl.

The UPLC-MS/MS system consisted of a UPLC (ABSciex, Shimadzu) attached to ABSciex 6500+ QTRAP mass spectrometer with ESI source. The Acquity BEH C18 (2.1mm x 50mm, 1.7μm) (Waters, Milford, MA, USA) column was used for the separation of compounds, and column oven temperature was +40°C. The autosampler temperature was set to +25°C. Injection volume was 2 μl. The mobile phases were water (A) and acetonitrile (B) both with 0.1% of formic acid and the flow rate was 0.6 ml/min. The linear gradient started from 30% B and proceeded to 98% in 6.5 min, followed by 1.5 min at 98% B, giving a total run time of 8 min. The data was normalized to dry weight (DW) and to the peak area of internal standard. The Analyst software (ABSciex) was used for data processing and quantification.

PTSI derivatization reagent generated betulin p-toluenesulfonyl carbamic diester, betulinic acid p-toluenesulfonyl carbamic ester, and testosterone toluenesulfonyl carbamic ester. Two MRM transitions were selected for each analyte, one for quantification and other for qualification. The ratio between quantification (quan) and qualification (qual) transitions should stay stable among runs. The transitions were as follows: betulin MRM 835.3 → 620.3 quan [M-PTSI-H_2_O-H]^−^, 835.3 → 638.3 qual [M-PTSI-H]^−^, betulinic acid MRM 652.3 → 455.2 quan [M-PTSI-H]^−^ 652.3 → 437.2 qual [M-PTSI-H_2_O-H]^−^, and testosterone 484.2 → 287.2 quan [PTSI-H]^−^, 484.2 → 269.2 qual [PTSI-H_2_O-H]^−^. ESI source temperature was set to 450 °C. ESI/MS/MS detection was performed in negative ion mode with ion spray (IS) voltage of −4000, curtain gas (CUR) 30, collision gas (CAD) at medium, entrance potential (EP) −10, declustering potential (DP) −60 (betulinic acid, testosterone) or −100 (betulin), collision energy (CE) −50, collision cell exit potential (CXP) −10.

### Cloning and mass spectrometry of lupeol synthase and CYP450 monooxygenase enzymes

Two major cloning constructs were designed for betulin biosynthesis using pRS424 vector (Burgers, 1999). This version of pRS424 vector contained two multiple cloning sites (MCS), one under GAL1 promoter and the second under GAL10. First, single constructs of CYP450 monooxygenase enzymes were isolated from both *B. pendula* (pRS424::CYP716), as well as four homologous CYP450 monooxygenases from *I. obliquus* (pRS424::CYP450). The second construct was a double insertion (pRS424::LUS-CYP716) of lupeol synthase (Bpev01.c0219.g0020.m0001 under GAL10 promoter) and CYP450 monooxygenase (Bpev01.c0219.g0021.m0001, under GAL1 promoter) enzymes isolated from *B. pendula* in pRS424 vector. All the vector constructs were transformed to yeast strain (*Saccharomyces cerevisiae* [w303 background]). The transgenic yeasts were grown and induced according to Zhou et al. (Zhou, Li, Li, & Zhang, 2016), with minor changes. SD-TRP was us as the drop out medium. For single inserted vectors, we used 50um lupeol (dissolved in DMSO:EtOh [1:1]) in induced growth media. After 60 hours of induction, the yeast growth media were centrifuged, and both media and the cell pallets were collected and sent for mass spectrometry.

Total of eight samples (yeast cells (4 tubes), and cell culture media (4 tubes)) were analyzed with UPLC-QTRAP/MS (MRM). Three triterpenoids (betulin [BE]) were extracted first from the media twice with 1.0 ml ethyl acetate (Merck) for 60 min in RT and centrifuged for 5 min in 15 000 rpm according to Cao et al. (G. Zhao et al., 2007). Testosterone was used as an internal standard (ISTD, 1.0 μl, 1.0 μg/ml). The cells were extracted in a similar manner as media, but yeast cells with 500 μl H_2_O and 1000 μl chloroform twice and were disrupted with freeze/thaw cycle (3 cycles) with ultra-sonication (15min) prior to extraction procedure.

The upper ethyl acetate was evaporated to dryness with MiVac Duo concentrator +40°C (GeneVac Ltd., Ipswich, UK). The residue was re-solubilized in 100 μl ACN. Due to very low concentration of lupeol, betulin and betulinic acid, extracts had to be derivatized with p-toluenesulfonyl isocyanate (PTSI) (Hu et al., 2013; Zuo et al., 2005) to improve sensitivity. Samples were derivatized in RT for 3 min with 10 μl of 60% p-toluenesulfonyl isocyanate (PTSI) (Sigma Aldrich) in ACN (Hu et al., 2013; Zuo et al., 2005). The derivatization reaction was terminated with 90 μl of MeOH with 30 s vortex mixing, giving the total volume of 200μl. Immediately after PTSI-derivatization, the MRM analysis of lupeol, betulin and betulinic acid was executed with UPLC-QTRAP/MS (ABSciex).

The UPLC-MS/MS system consisted of ABSciex UPLC attached to ABSciex 6500+ QTRAP mass spectrometer. The separation of the analytes column was Acquity BEH C18 (2.1mm x 50mm, 1.7μm) (Waters, Milford, MA, USA), with the temperature of +40°C. The autosampler temperature was set to +25°C. The injection volume was 10μl. The chromatographic conditions were executed as described previously at Hua *et al.* (Hu et al., 2013). The mobile phase consisted of water with 0.1% of formic acid in H_2_O (A) and acetonitrile (B) with a flow rate of 0.6 ml/min. The linear gradient started 30% B and proceeded to 98% in 6.5 min, left in 98% B for 2 min, and switched back to initial conditions and left to stabilize, giving a total analysis time of 10 min.

ESI source temperature was set to 450°C. ESI/MS/MS detection was performed in negative ion mode with ion spray (IS) voltage of −4000, curtain gas (CUR) 30, collision gas (CAD) at medium, entrance potential (EP) −10, de-clustering potential (DP) −60 (betulinic acid, testosterone) or −100 (betulin), collision energy (CE) −50, collision cell exit potential (CXP) −10. The Analyst software (ABSciex) was used for data processing. PTSI derivatization reagent generated betulin p-toluenesulfonyl carbamic diester (BTCD). The transitions for betulin and ISTD (testosterone) was as follows: Betulin (BE) MRM 835.2 → 620.2 [M-PTSI-H_2_O-H]^−^, 835.2 → 638.3 [M-PTSI-H]^−^ and 835.2 → 196.0 [PTSI-H]^−^, and for testosterone MRM 484.2 → 287.2 [M-PTSI-H]^−^ and MRM 484.2 → 269.2 [M-PTSI-H_2_O-H]^−^. The most intense transitions of MRM 835.2 → 620.2 (betulin), and MRM 484.2 →287.2 (ISTD, testosterone) were used.

## Results and discussion

### Nuclear and mitochondrial genome assemblies and annotations

Pacbio sequencing of *I. obliquus* strain from Merikarvia yielded 4.82 Gb of data (96x coverage) with N50 read length of 9,200 bp. Falcon assembly resulted in a 41.1 million bp genome, consisting of 301 primary contigs with an N50 value of 516 kilobases. Overall, the genome size of *I. obliquus* was comparable with other species from *Hymenochaetales* (Supp. table 1).

Genome annotation of primary assembly yielded 13,778 gene models with 91.7% of universally conserved single-copy genes being present (BUSCO v3.0, fungi database) (Waterhouse et al., 2017). To support gene model prediction, RNA-seq was carried out from total RNA extracted from *I. obliquus* reference strain sample grown on wood dust. Genome-guided *de novo* assembly of RNAseq data showed 91.3% BUSCO completeness with 31% duplicated genes (Supp. table 1). Altogether 70.8% of *I. obliquus* genome consisted of coding sequence, with 53.1% in exons. Mean intron, exon, CDS and gene lengths were 89, 284, 1447, and 2113 base pairs, respectively (Supp. table 1). Mitochondrial genome was also assembled and annotated as described previously (Salojärvi et al., 2017), resulting in 29 tRNAs, 32 coding sequences, and 3 rRNAs.

Transposable elements (TEs) have been suggested to play a major role in genome plasticity and evolution. Thus, the classification and characterization of genes in close proximity of TEs are of general interest, especially in the case of pathogenic organisms (Faino et al., 2016). In *I. obliquus*, the total genome repeat content was found to be 26%, with 14.24% of repeats being unclassified. The percentage of retrotransposon elements was 8.37%, and DNA transposon elements 1.2%. In contrast to retrotransposon elements, *I. obliquus* genome contained higher amounts of DNA transposon elements compared to the related *F. betulina* and *F. mediterranea* (Supp. table 1). Unlike *F. mediterranea* (42.27% repeat sequences), TE content of *I. obliquus* did not fully explain its large genome size (26% of repeat sequences) (Hage et al., 2021). Gene models flanking the upstream and downstream of TEs contained mainly transposition elements, and gene clusters between two transposable elements from the same DNA transposon class suggested the enrichment of gene models involved in transmembrane transporter (GO:0003677), protein dimerization (GO:0046915), transposition (GO:0046983 and GO:0006310), and DNA biding and recombination (GO:0006313 and GO:0032196) (Supp. table 1) (Ali et al., 2014; Kang, Lebrun, Farrall, & Valent, 2001). The enrichment of the last two categories suggests that some of the predicted gene models may be unidentified transposable elements, and they are organized as clusters in the genome.

### Secreted proteins in *I. obliquus*

The exact mechanisms of *I. obliquus* pathogenicity and its modes of interaction with the host are not known, but characterization of secreted proteins is the first step to shed more light on the mechanisms involving the initial penetration of plant defences by effector proteins. Secreted proteins (SPs) are known for their essential role in pathogen-host interactions. Altogether 1052 open reading frames (ORFs), 7.6% of all gene models, were predicted as possible secreted proteins, with minimal known homologs (Supp. table 2). The SPs were scattered across 128 contigs. Most of the ORFs were likely species-specific, since homology searches with known secreted proteins from other species were successful for only 110 of the ORFs (Supp. table 2). Twenty-one ORFs overlapped at least with one class of TEs, having eighteen unclassified categories. Total of 988 ORFs were co-localized between two TEs of the same TE class, suggesting a role for TEs in SP evolution and diversification.

### Carbohydrate-active enzymes

The palette of carbohydrate-active enzymes (CAZymes) present in the genome dictate to a large extent the modes of substrate utilization by the fungus (Eastwood et al., 2011; Navarro et al., 2021). We identified 466 candidate genes classified as carbohydrate-active enzymes (CAZymes), known for their biological roles in anabolism and catabolism of different carbohydrates such as glycogen, trehalose, and glycoconjugates (Supp. table 5). Altogether 211 enzymes were classified as glycoside hydrolases (GHs) and 43 were categorized as carbohydrate binding modules (CBMs) whereas the overall number of glycosyltransferases (GTs) and carbohydrate esterases (CEs) were found to be 110 and 23, respectively. Finally, 10 enzymes were assigned to polysaccharide lyases (PLs) (Supp. table 5). Overall the CAZyme palette was similar to other lignin-degrading fungi (Liu et al., 2019). RNAseq analysis of *I. obliquus* grown on *B. pendula* wood dust and publicly available RNAseq data (Fradj et al., 2019) identified in 214 (out of 466) CAZymes with positive Log_2_FC (Supp. table 5) in at least one of the experimental conditions. Majority of DE CAZymes belonged to glycoside hydrolases and glycosyltransferases categories.

### Phylogenomics and expanded gene families in I. obliquus genome

In order to estimate a taxonomic placement for *I. obliquus*, we collected the proteomes of fifteen representative fungal species from different orders among *Basidiomycota*: all sequenced species within *Hymenochaetales* (*Fomitiporia mediterranea, Inonotus obliquus, Onnia scaura, Phellinidium ferrugineofuscum, Porodaedalea niemelaei, Trichaptum abietinum, Rickenella mellea, Schizopora paradoxa),* and representatives of *Russulales (Heterobasidion annosum)*, *Polyporales (Fomitopsis betulina, Postia placenta, Phanerochaete chrysosporium)* and Agaricales (*Laccaria bicolor, Coprinopsis cinerea, Schizophyllum commune)*. To root the taxonomy we added five outgroup species, including two representatives of the other major classes in *Basidiomycota*: *Pucciniales (Puccinia graminis)* and *Ustilaginales (Ustilago maydis*), as well as three model *Ascomycota* species from *Saccharomycetales (Saccharomyces cerevisiae)*, *Schizosaccharomycetales (Schizosaccharomyces pombe),* and *Sordariales* (*Neurospora crassa)*. We next clustered the full proteomes into gene families (orthogroups) using Orthofinder and identified single copy orthogroups. Phylogeny estimation was carried out using 4040 single copy gene groups and rooted to *Ascomycota* species. The resulting tree illustrates the known taxonomy among the 20 fungal species (Figure 1; Supp. table 3) and the split of *Hymenochaetaceae* family occurs at the expected phylogenetic position (Hibbett & Thorn, 2001; Matheny et al., 2007; R.-L. Zhao et al., 2017) (Figure 1). Interestingly *R. mellea* was placed together with the *Russulales* representative, further analysis is however beyond the scope of the present work.

**Figure 1.**
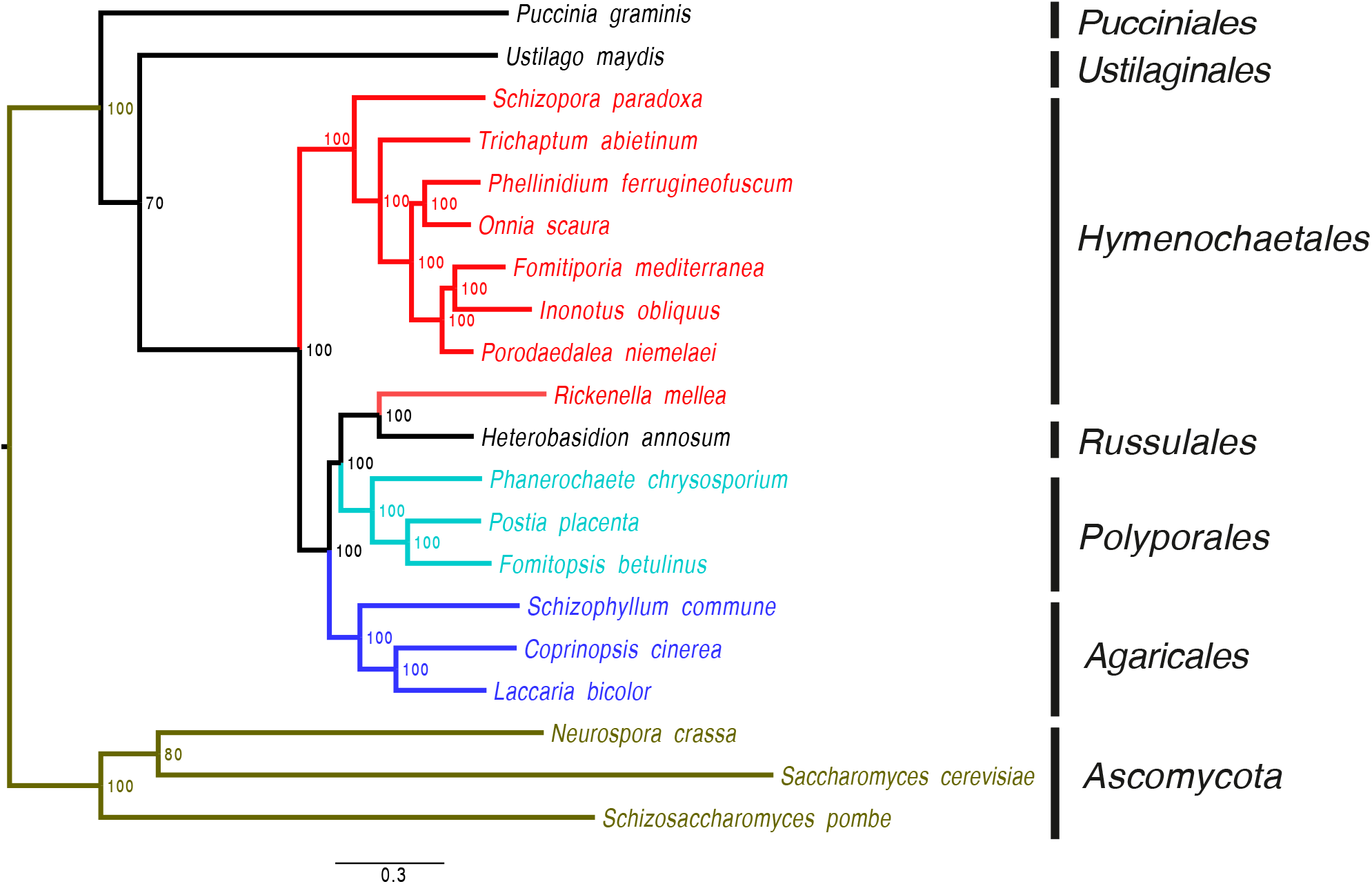
Phylogenetic tree of three *Ascomycetes* and 17 *Basidiomycetes* species. *Ascomycetes* are highlighted with green colour and grouped by Phylum. *Basidiomycetes* species are grouped by taxonomic order. Bootstrap values are plotted next to the nodes illustrating the level of the confidence of the split, and the phylogenetic tree was rooted to *Ascomycetes* clade.

To look for gene family evolution we then identified gene families that were expanded in *I. obliquus*. Altogether 167 orthologous gene clusters were significantly expanded in comparison to the other nineteen fungal species (chi-squared test; Supp. table 4**Error! Reference source not found.**). The expanded gene families were enriched for 23 GO terms such as terpene synthase (GO:0010333), oxidoreductase (GO:0016684), and hydrolase (GO:0016788) activities. In addition, GOs related to oxidative stress responses (GO:0006979), transposition (GO:0015074, GO:0006313), and protein dimerization activity (GO:0046983) were also expanded (Supp. table 4); most of these categories involve members of cytochrome P450 gene family.

### Genome evolution in *I. obliquus*

The high-quality whole genome assembly allowed us to gain further insight into the gene family evolution by synteny analyses using self-self alignments. The analysis suggested a recent whole genome duplication (WGD) event in *I. obliquus*. Based on the synonymous mutation (Ks) spectrum the event occurred after the split from *F. mediterranea* (approximately 112 million years ago during the Triassic period; (Kumar, Stecher, Suleski, & Hedges, 2017)). Similarly, an independent lineage-specific WGD was observed also in *P. niemelaei* (Figure 2).

**Figure 2.**
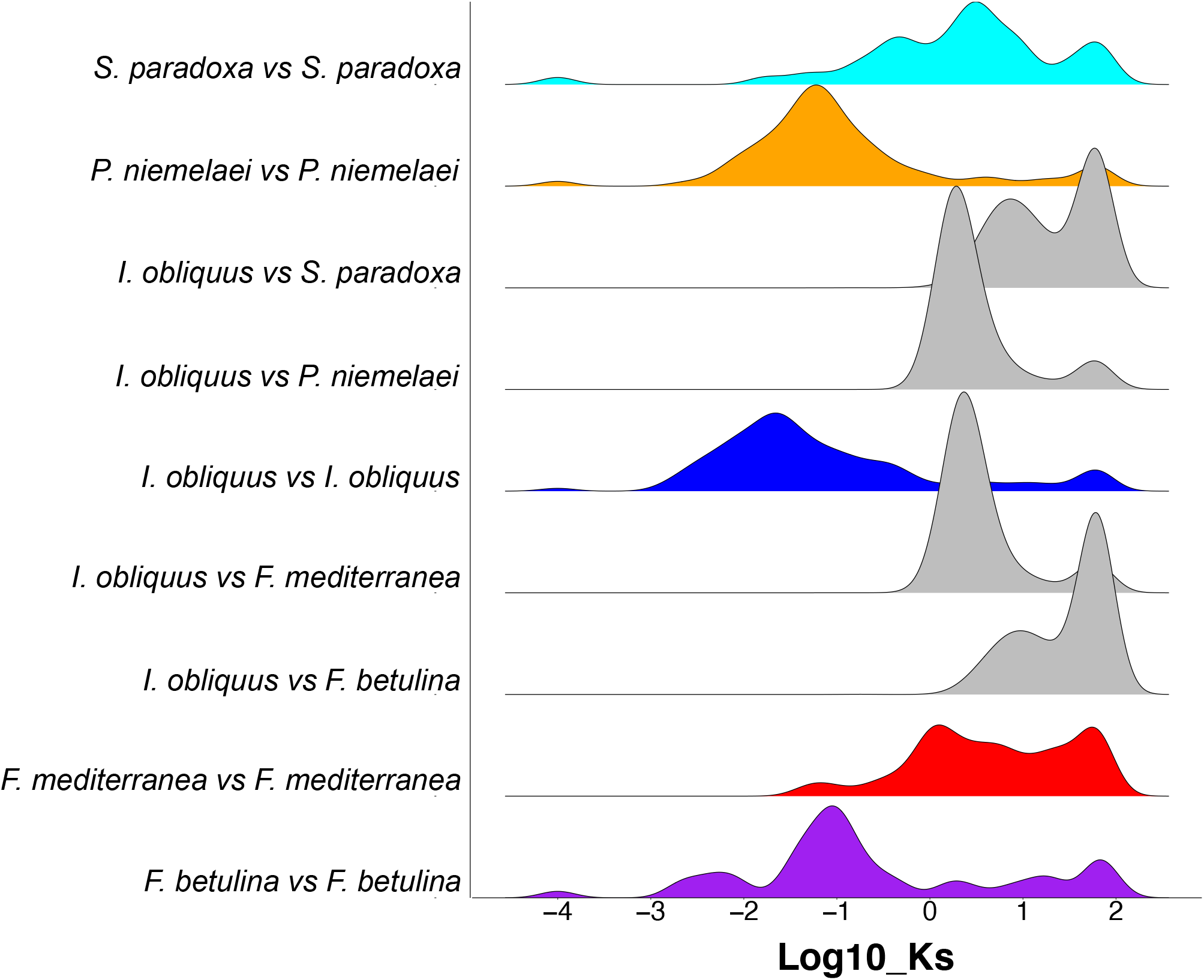
Density plots of the number of synonymous (Ks) substitutions in syntelogs identified from syntenic alignments of *I. obliquus*, *F. mediterranea*, *P. niemelaei*, and *S. paradoxa*. X-axis is displayed as log10 of synonymous substitutions per synonymous site (Ks).

Synteny analysis identified a total of 1,112 genes originating from the whole genome duplication event, whereas a considerably higher amount, 6,200 genes, were identified in tandem duplications (Supp. table 3). The tandemly duplicated genes reflect the shorter-term adaptation in the species, and in general have been found to be associated with environmental responses (Panchy, Lehti-Shiu, & Shiu, 2016). In *I. obliquus*, the tandemly duplicated genes were enriched for carbohydrate biosynthesis, heme binding, oxidoreductase activity, tetrapyrrole binding, and DNA transposition. In contrast, syntenic regions harbored genes related to biological pathways such as terpene synthesis and cell cycle (Figure 3, Supp. table 6). The overlaps between tandemly duplicated, syntenic genes and expanded gene families were significant (p-value=6.1094e-11, Fisher exact test) (Figure 4). Altogether, the genome evolution analyses highlight the significance of tandem duplication events in adaptation of the *I. obliquus* to different ecological niches and the central role of secondary metabolism and particularly the expansion of CYP450 gene family by small scale duplication events (Figure 5).

**Figure 3.**
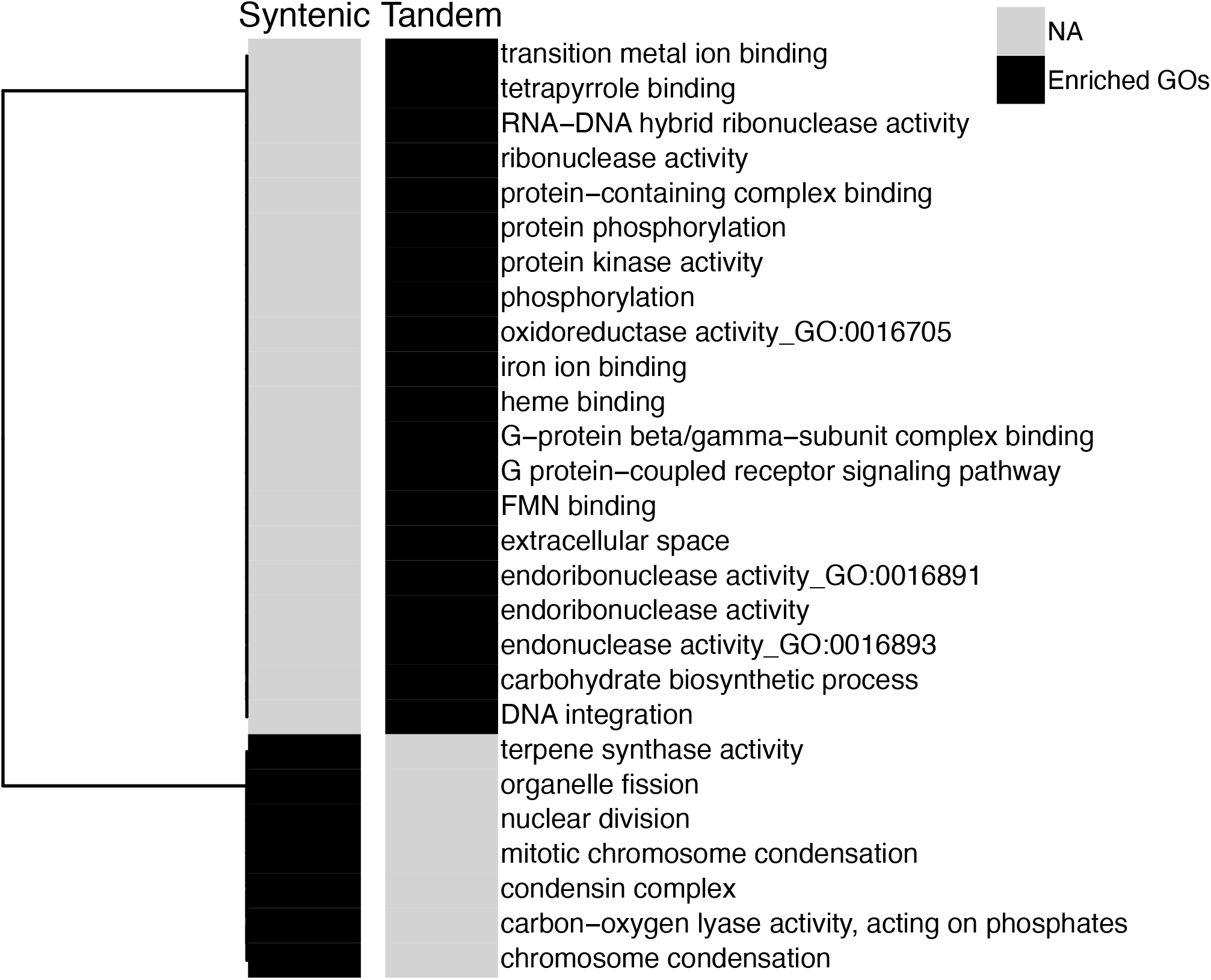
Gene ontology (GO) enrichment of genes identified in syntenic and tandemly duplicated regions in *I. obliquus* and four other species. Separate heatmaps are shown for the syntenic and tandemly duplicated regions, black color shows significantly enriched GOs and grey illustrates non-significant enrichments (NS). GOs are clustered using the Euclidean distance and hierarchical clustering (method: complete).

**Figure 4:**
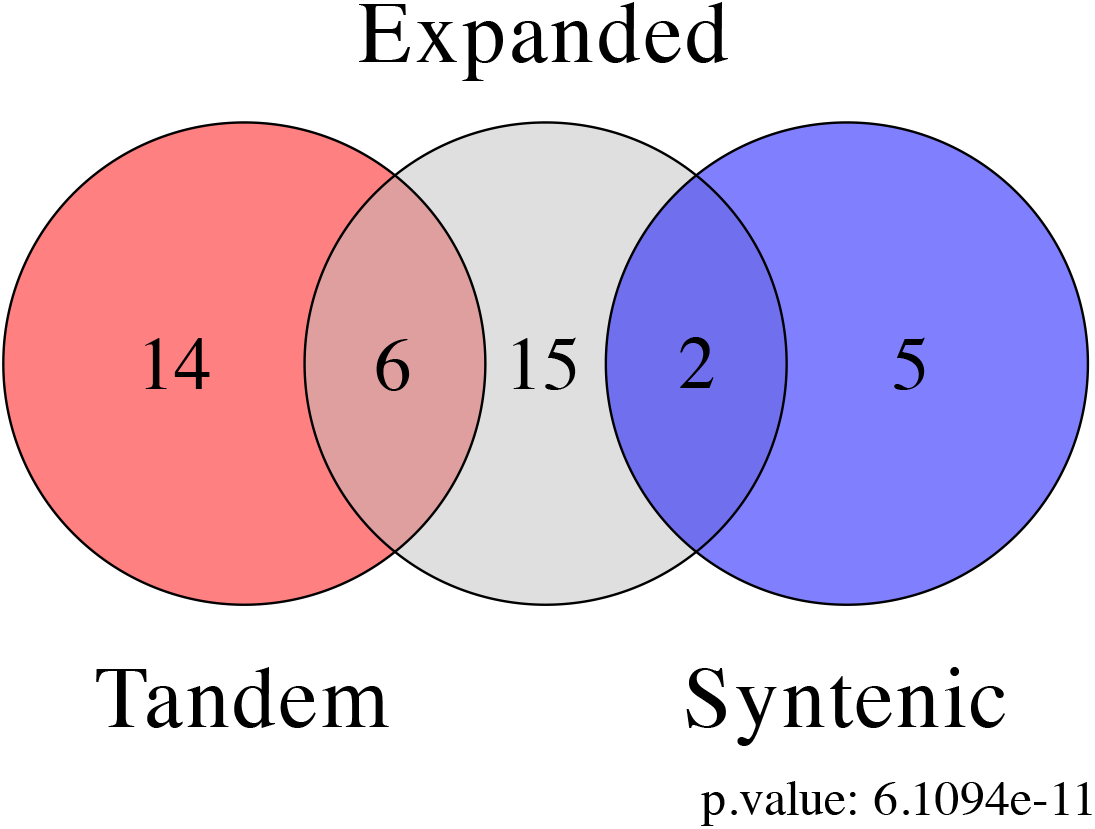
Venn diagram of GO enrichment analysis for expanded gene families and tandemly duplicated genes from *I. obliquus* genome. Each category is highlighted and labeled by specific color. P-value was calculated with Fisher exact test assessing the the statistical significance of the overlap. Tandem: tandemly duplication genes; Expanded: expanded gene families; syntenic: syntenic self-self alignment of *I. obliquus* resulting in the set of genes originating from whole genome duplication event.

**Figure 5.**
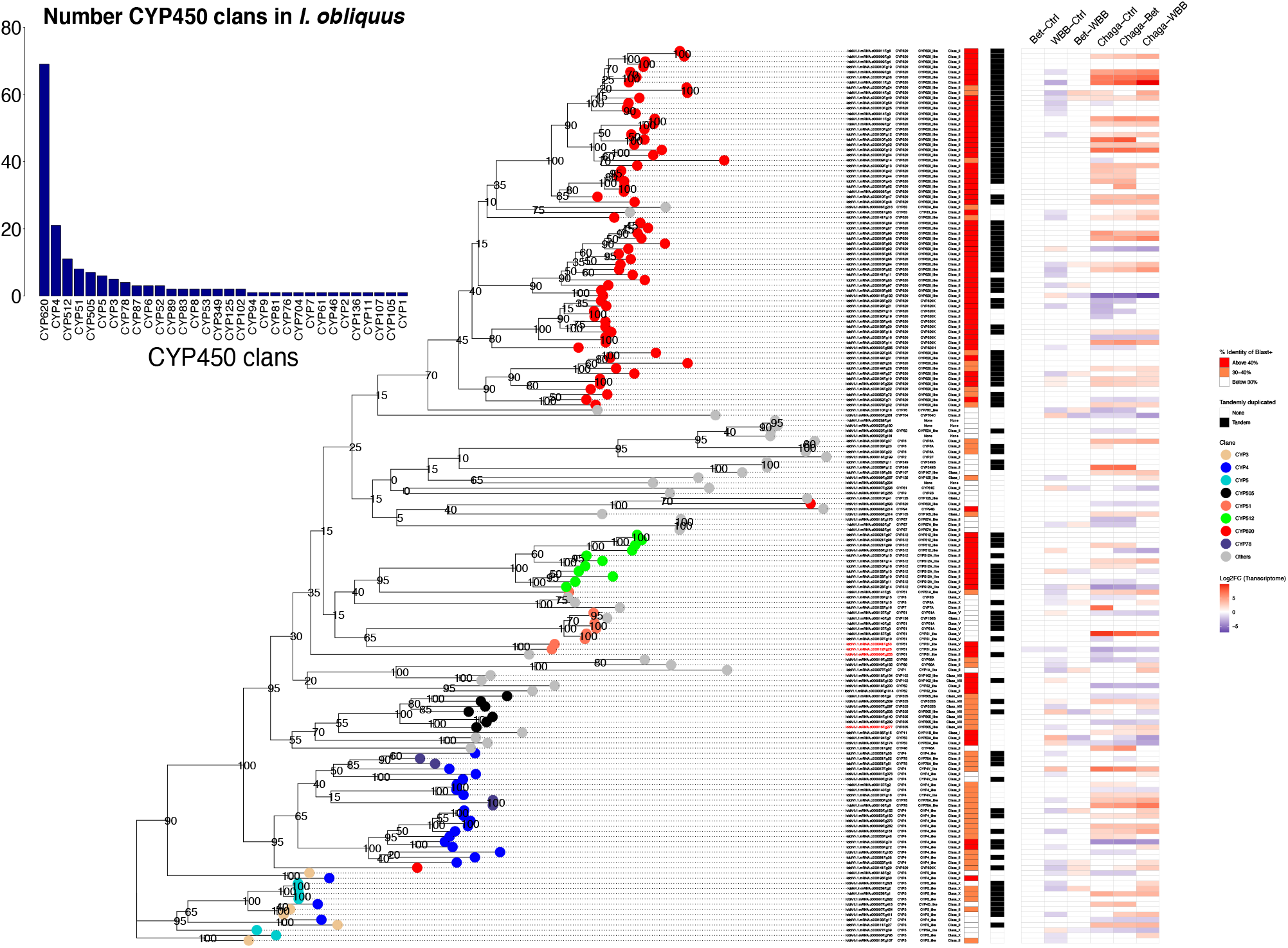
A gene tree of CYP450s predicted in *I. obliquus*. The clades are highlighted according to the major clan of the enzyme and tandemly duplicated genes are indicated with black squares; the color scheme is shown in the color key to the right of the plots. The branches are labeled by gene ID, cytochrome P450 clans, family, and class. The heatmaps illustrate BLAST similarities (red), tandemly duplicated (black/white), and differential expression of the genes. Barplot in the inset shows the number of CYP450 clans in *I. obliquus*.

The members of CYP450 family have critical roles in fungal metabolism and adaptation to specific ecological niches. Altogether 172 CYP450 monooxygenases were predicted in chaga, suggesting a complex biochemical diversity in chaga metabolism. Division into clans and families revealed that most of the enzymes belonged to clan CYP620 (69 members), followed by CYP4 and CYP512 clans (Figure 5). CYP620 is shown to be involved in terpenoid synthases (Yap et al., 2014; Yu, Song, Liang, Wang, & Lu, 2020). CYP4 is studied predominantly in phylum of *Arthropoda*, and shown to be involved in biosynthesis of endogenous compounds (Zhu, Moural, Shah, & Palli, 2013). Finally, CYP512 clan has been hypothesised to have catalytic activities towards steroidal-like compounds, primarily testosterone (Ide, Ichinose, & Wariishi, 2012).

A high proportion CYP450 gene models, 79 out of the total of 172, were tandemly duplicated, and many were members of CYP620 clan. GO enrichment analysis of two genes upstream and two genes in downstream of all CYP450 monooxygenase enzymes suggested the enrichment of biological functions related to oxidoreductase activity, heme binding, transmembrane transporter activities, and tetrapyrrole binding (Supp. table 6). These results suggest tandem duplications of CYP450s, and additionally the colocalization with cytochrome P450 reductase (CPR) partners, facilitates their functional divergence (Ebrecht et al., 2019).

The observed colocalization of genes related to CYP450s suggested the presence of biosynthetic clusters in the *I. obliquus* genome. We therefore sought biosynthetic gene clusters by antiSMASH (Blin et al., 2013), identifying altogether 24 clusters in 17 contigs: 15 terpene synthase, 3 polyketide synthase, and 4 non-ribosomal peptide synthetase clusters. The clusters were significantly enriched for tandemly duplicated genes (Fisher exact test, p.value=6.34E-76), suggesting that tandem duplications are a dominant process in their diversification. Furthermore, the clusters were enriched for CYP450 gene family (p-value: 0.03161975), highlighting their central enzymatic role in secondary metabolism (Supp. table 6).

### Metabolomics fingerprinting of terpenoid compounds in five *I. obliquus* strains and *F. mediterranea*

Since chaga showed a significant expansion of CYP450 genes and a considerable number of biosynthetic clusters, we next carried out metabolic fingerprinting of *I. obliquus* to study whether the secondary metabolism was indeed diversified in chaga compared to *F. mediterranea*. Terpenoid fingerprints in five strains of *I. obliquus* were distinctly different from *F. mediterranea* (Supp. table 1). Altogether the chaga strains showed 546 mass spectrum peaks, and only 135 of them were shared with *F. mediterranea* (Figure 6 A). In addition, pairwise comparisons of each chaga strain and *F. mediterranea* found 178 metabolomic features among *I. obliquus* strains with significantly higher abundance (Supp. Fig 2, Supp. table 7). Many of the peaks were predicted to have molecular formulae with 30, 31, and 28 carbon backbones, similar to lupeol, betulin and betulinic acid. Among *I. obliquus* strains, Merikarvia had distinct metabolomic fingerprints and clustered more distant from other strains (Figure 6 B, C) in principal coordinate analysis. This suggests that genotypic variation plays a role in metabolic diversity (Figure 6 C). With regards to betulinate compounds, Merikarvia strain had higher abundance of betulin and betulinic acid production in comparison to other strains of *I. obliquus*. There were no peaks which resembled the standard for betulin or betulinic acid (98%, Sigma-aldrich) in *F. mediterranea*, but a significant quantity of lupeol-like substance was discovered (Supp. Fig 2).

**Figure 6:**
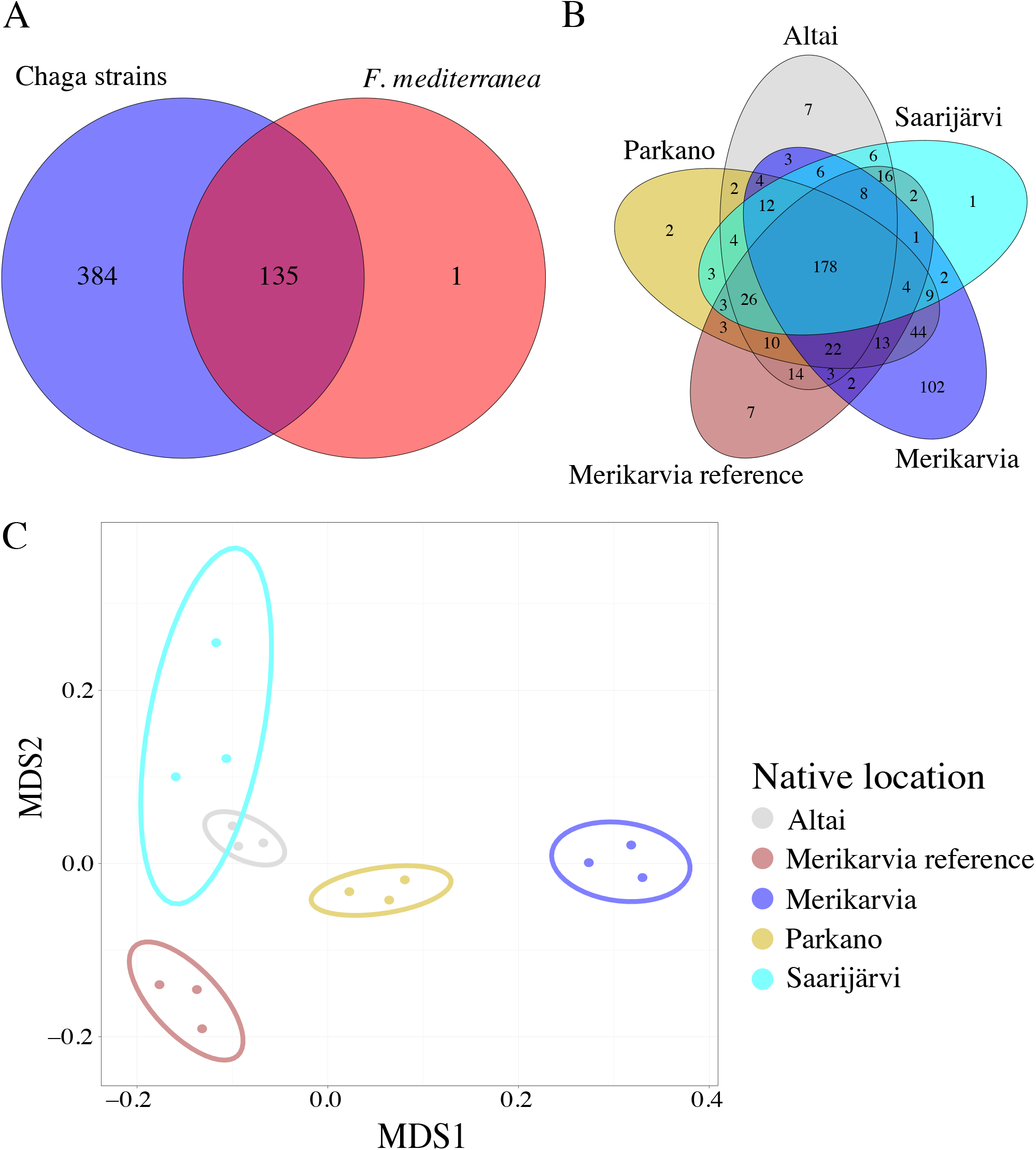
Venn diagr ams of UPL C-MS mass spectrum for metabolomic fingerprints. A) Pooled mass spectrums from five strains of *I. obliquus* and one *F. mediterranea,* B) mass spectrums of five strains of *I. obliquus*, C) multidimensional scaling (MDS) plot of metabolomics abundant of five strains of *I. obliquus*. Panels B and C use the same color coding, explained in panel C legend. .

### Functional analysis of lupeol synthase and CYP450 monooxygenase

Even though metabolic fingerprinting does not identify the underlying metabolites, the analysis suggested the presence of terpene and lupeol as well as betulin derivatives based on the predicted carbon backbones. Intraspecies quantification of betulin and betulinic acid (Using HPL) among six species of betula and three strains of chaga showed a higher concentration of betulinic acid compared to betulin in chaga strains. In contrast, the opposite result was observed in six species of Betula, where the concentration of betulin was consistently higher in comparison to betulinic acid (Figure 7).

**Figure 7:**
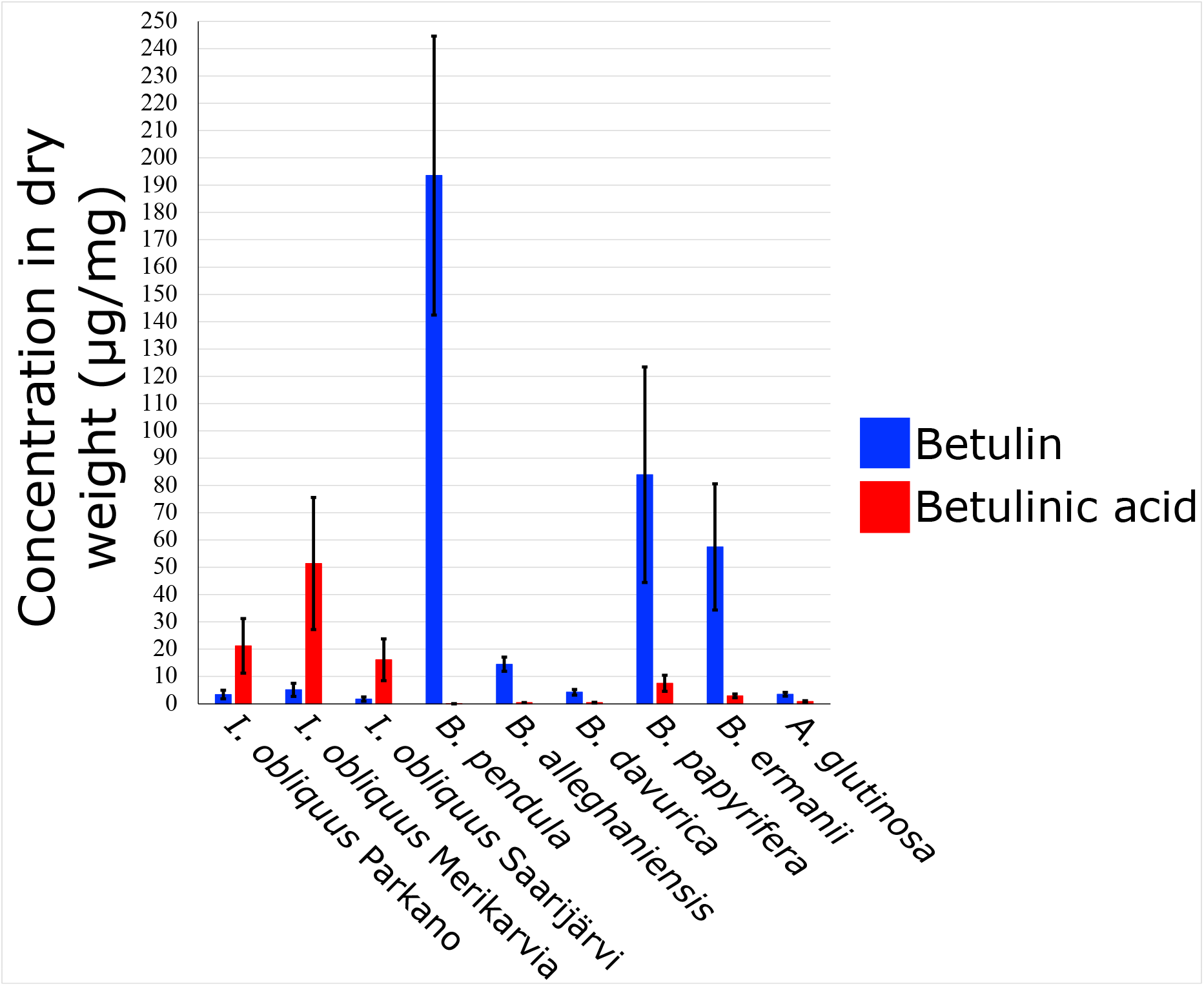
Concentration of betulin and betulinic acid in three strains of chaga and six betula species by HPLC-MS. Betulin and betulinic acid are highlighted in red and blue.

The high quality gene model predictions allowed us to look for the candidate enzymes responsible for the betulin biosynthesis in chaga. Since no members of CYP716 family were not predicted in chaga we identified four best candidates based on homology analysis of CYP450s to known CYP716 family members from plant species, such as *B. pendula*. Yeast expression system has been used successfully for cloning CYP450 monooxygenase enzyme from *B. platyphylla* (Zhou et al., 2016). To functionally validate our candidate genes, we first constructed single insert expression vector for birch CYP716 enzyme (pRS424::CYP716). In addition, we also constructed a double insert vector where lupeol synthase and lupeol monooxygenase from *B. pendula* (Safronov et al., 2019) were inserted into two multiple cloning sites of a vector (pRS424::LUS-CYP716), which was then transferred into yeast expression system. Similar to single insert vector for CYP450 monooxygenase enzyme from *B. pendula*, we isolated four CYP450s from *I. obliquus* and cloned them to engineer single inserted constructs. The single constructs were grown in media which was spiked with standard lupeol compound (98%, Cayman) as the precursor. In contrast to single insert vectors, double insert vector from *B. pendula* (pRS424::LUS-CYP716) expressed lower concentration of betulin compared to single insert vector (pRS424::CYP716) from *B. pendula*. These differences might be explained by the lower initial amount of available precursor compound, lupeol, for CYP716 monooxygenase (Figure 8). In addition to *B. pendula* constructs, all four candidate CYP450 monooxygenases from *I. obliquus* (pRS424::CYP450) showed some degree of betulin production when compared to standard betulin (98%, Sigma-Aldrich) spectrum. Among the four candidates, the enzyme with gene ID c000016F_g277 (clan CYP505) had the highest amount of betulin production. Interestingly, the cDNA length of this enzyme was 3297 bp, almost twice the length of the other three candidate CYP450 monooxygenase homologs from *I. obliquus*. Upon close examination the amino acid and nucleotide sequences of c000016F_g277 resemble a chimeric isoform of two CYP450 monooxygenase enzymes (Figure 8). In general, our study showed that yeast cell fractions contained higher concentration of betulin compared to the culture media fractions (Figure 8), thus confirming the function of the inserted enzymes.

**Figure 8:**
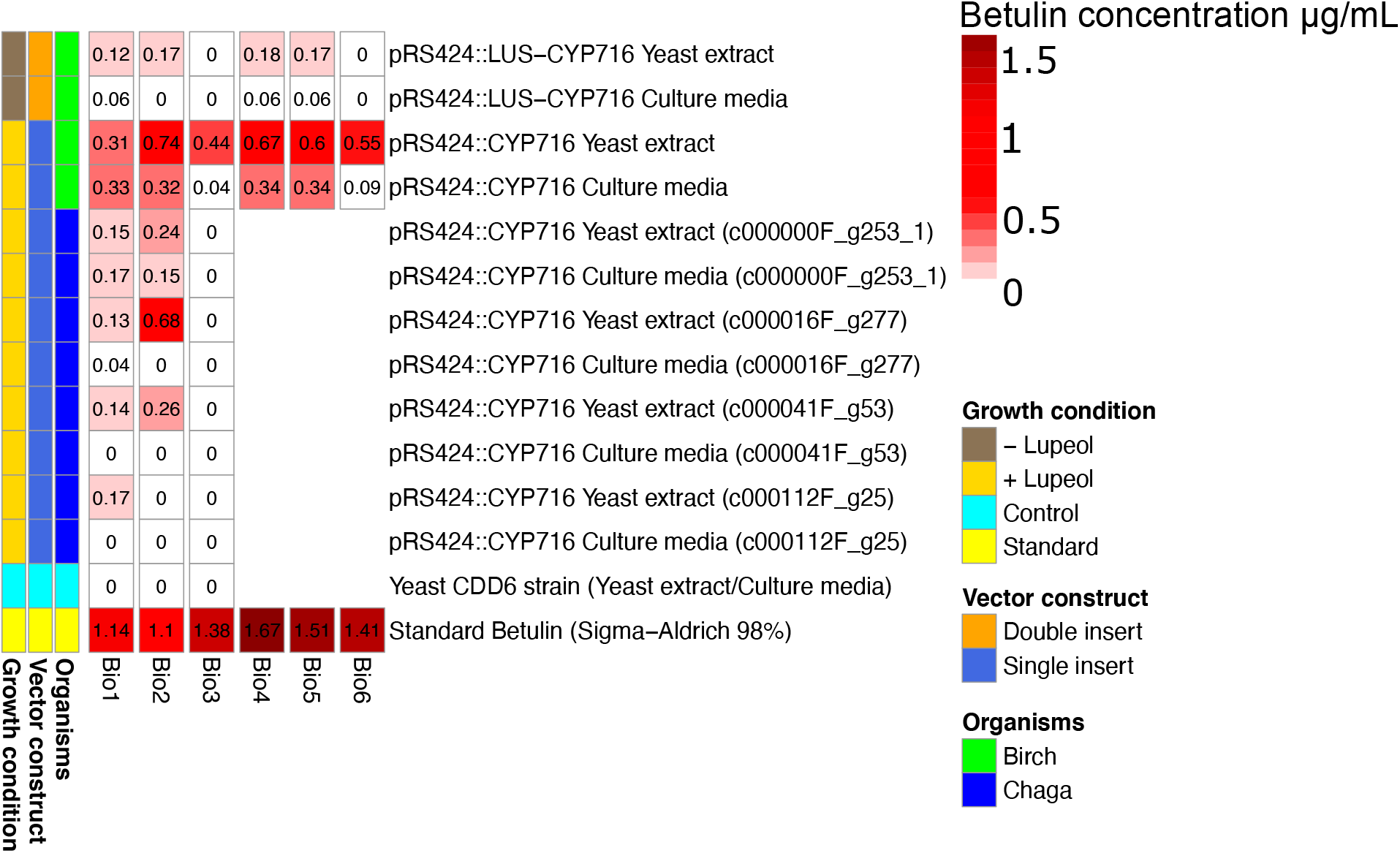
Heatmap illustrating the amount of betulin synthesized in transgenic yeast. The columns illustrate the biological replicates, and rows the vector constructs for lupeol synthase and CYP716 genes. The biological source for lupeol synthase was birch (green, *B. pendula*), and the biological sources for CYP716 enzymes were from birch and chaga mushroom (dark blue, *I. obliquus*). Vector constructs were divided to double insert (orange, lupeol synthase and CYP716 from birch in a single vector), and single inserts (royal blue, only CYP716 from birch and chaga). The color palette of the heatmap illustrates the concentrations (μg/mL) of betulin found in yeast cell (Yeast extract) and yeast growth media (Culture media), with white color assigned to minimum and dark red to maximum concentration of betulin. The second heatmap illustrates the growth conditions: brown (− Lupeol) is yeast growth media without standard lupeol (precursor for CYP716 gene), gold (+ Lupeol) yeast growth media with standard lupeol, and cyan CDD6 yeast strain used as control, and finally yellow is lupeol standard (98% purity).

To study the evolution of potential homologs and orthologs for the four candidate genes from *I. obliquus*, we carried out microsynteny analysis for the cloned CYP450 monooxygenase enzymes against four *Hymenochaetales* and *F. betulina* species. Orthologous one-to-one relationship with other fungal species was confirmed for c000112F.g25 (clan CYP51) and c000041F.g53 (clan CYP51) enzymes (Supp. Fig 3-C,D), whereas microsynteny analysis of c000000F.g253 (clan CYP61) enzyme found a cluster of homologous genes in 5’ and 3’ of the c000000F.g253 in *I. obliquus* genome (Supp. Fig 3-B). The microsynteny of the chimeric c000016F.g277 linked to a putative ortholog in *F. mediterranea* with similar organization (gene_7933, clan CYP505) (Supp. Fig 3**Error! Reference source not found.**-A), whereas in other *Hymenochaetales* the syntenic analysis identified two separate CYP450s. This suggests that the fusion gene has arisen from a non-homologous recombination event in the common ancestor of *F. mediterranea* and *I. obliquus*, and after divergence of *P. niemelaei* where the CYP450s were still found separate. Both c000016F.g277 and gene_7933 contain two heme-binding domains, but gene_7933 from *F. mediterranea* has three oxygen-binding domains (with AGADTT/GGDDTG motifs) instead of two in *I. obliquus* (AGADTT). Therefore the fusion may have occurred also independently in chaga and *F. mediterranea* (Supp. table 8), or then involved a loss of the third oxygen-binding domain in chaga.

### Evolution of conserved domains and phylogeny reconstruction of cytochrome P450 monooxygenase

Since betulin and betulinic acid are antifungal substances and they are produced in the main natural host of *I. obliquus*, it is possible that the enzymes have been introduced into chaga or its ancestor via horizontal gene transfer, either directly from the host species or then via another species cohabiting with chaga. However, a phylogenetic tree of a set of monooxygenase enzymes (77 enzymes) from *I. obliquus* and *F. betulina* with sequence similarity to plant CYP716, as well as CYP716 enzymes of eight plant species known to produce betulinate compounds, shows a distinct divergence of fungal clades from the plant species. This result is consistent both in protein and DNA based phylogenetic trees, providing no evidence of gene transfer events (Figure 9).

**Figure 9:**
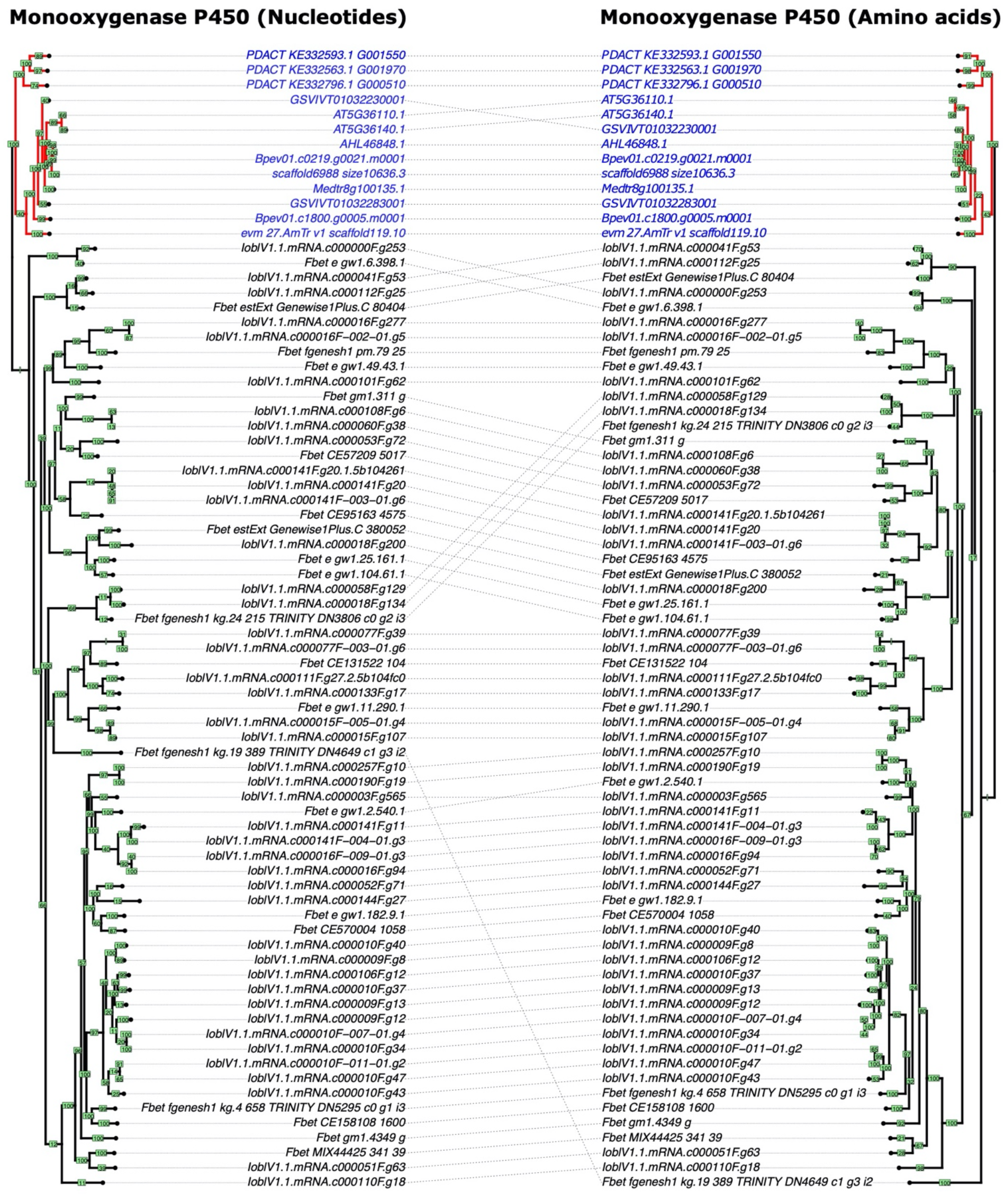
Phylogenetic reconstruction of 70 monooxygenase enzymes from *I. obliquus*, *F. betulina*, and 8 plant species. Trees are labeled according to the type of the sequence used in multiple sequence alignment (amino acid or nucleotide) and rooted to plant species. Cophylo (from phytools) function is used in order to rotate the branches to match the tips and the labels. Bootstrap values (green rectangular) illustrate the level of the confidence, and plant species are highlighted in blue.

**Figure 10:**
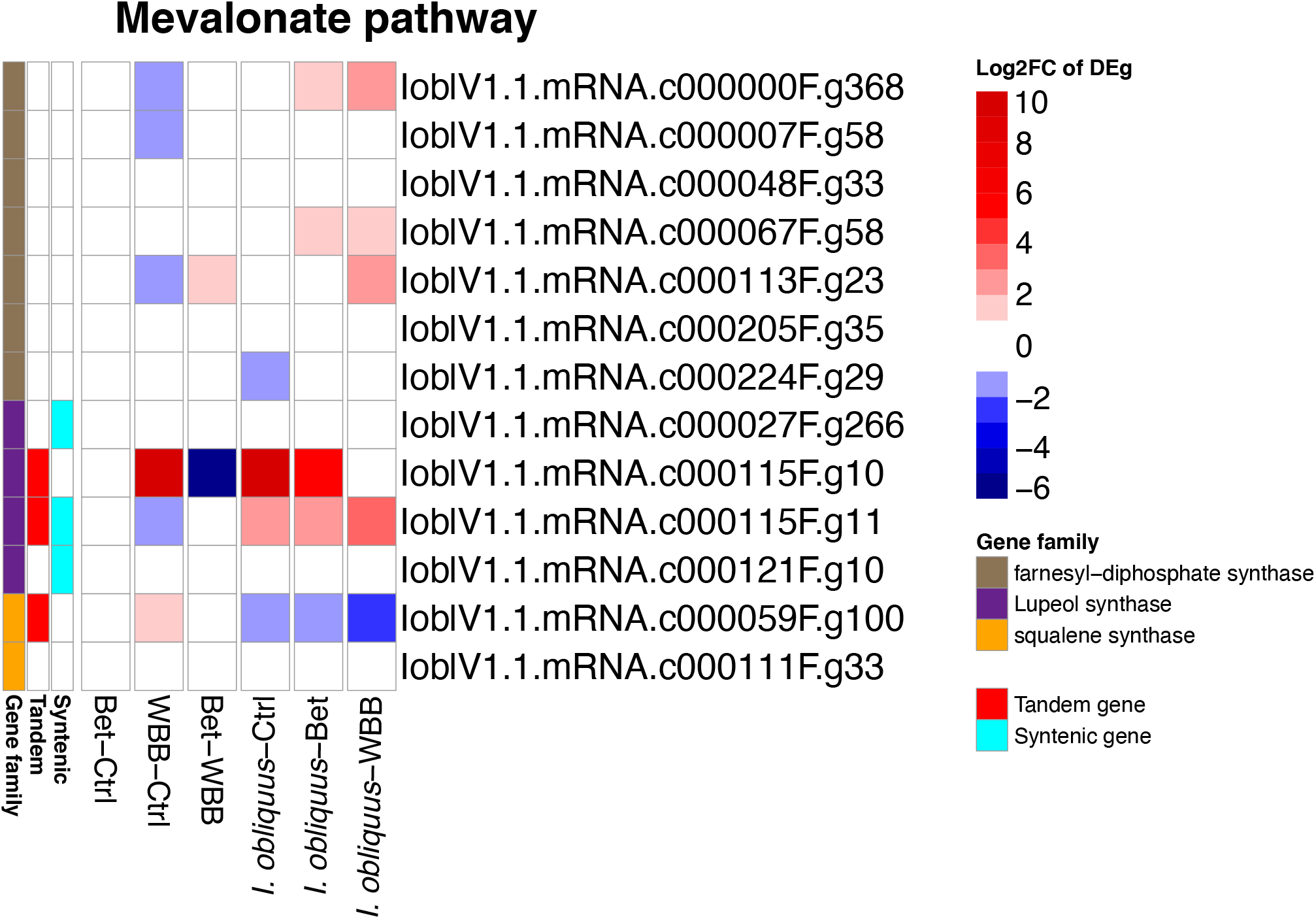
Heatmap of differentially expressed (DE) genes annotated: A) CYP450 monooxygenase enzymes, and B) key enzymes in mevalonate pathways (MVA). The larger heatmap illustrates the log_2_ fold-changes (log_2_FC) of DE genes, the color is proportional to differential expression. The smaller heatmap illustratres the duplication origins of the gene, either syntenic originating from whole genome duplication or tandem originating from segmental duplication. Genes are clustered using the Euclidean distance and hierarchical clustering (method: complete). Gene ID highlighted in blue was selected for cloning.

### Gene expression analysis

Total of 119 (out of 172) monooxygenase enzymes were significantly expressed with positive log_2_fold change (log_2_FC) values in at least one of the DEg comparisons (Figure 5) and three key enzymes involved in mevalonate pathways were among this set (**Error! Reference source not found.**). We also observed a pair of tandemly duplicated lupeol synthase enzymes to have the highest expression levels. The expression profiles of the chaga samples grown on *B. pendula* wood dust were stronger than the samples grown in culture media from (Fradj et al., 2019). When inspecting the expression profiles for genes with positive log_2_FC , enrichments were found for WD40-repeat binding (50 genes out of 294), melanin biosynthesis (65 genes out of 157), aquaporin (20 genes), lipases and peptidases (Supp. table 9).

## Conclusion

We observed genome evolution leading towards complex terpene biosynthesis in *I. obliquus*, both in genes originating from whole genome duplication events as well as tandem duplications within the CYP450 gene family. It is possible that the whole genome duplication event is associated with the initial expansion of terpenoid biosynthesis capacity in *I. obliquus*, since no such expansion was observed in the related species *F. mediterranea*. In contrast to eg plants, the number of whole genome duplication events in fungal kingdom has been low (Albertin & Marullo, 2012), but this may be due to faster genome evolution in fungi, making the WGDs difficult to identify (Campbell, Ganley, Gabaldón, & Cox, 2016). The CYP450 superfamily is associated with many reactions in secondary metabolism, and through metabolomics fingerprinting we confirmed that the fungus indeed produces a rich palette of terpenoid derivatives. However, we found no evidence of a horizontal gene transfer event between *B. pendula* and *I. obliquus*, and the identified candidate lupeol monooxygenases in *I. obliquus* were members of a different CYP505 clan with low sequence similarity to their birch counterpart enzymes. Therefore CYP450 monooxygenases enzymes responsible for betulinate biosynthesis in the two species most likely result from convergent evolution.

## Author’s contributions

O.S and J.S conceived and designed the project. Funding acquisition is carried by J.S and J.K. O.S collected the DNA and RNA samples. O.S and J.S managed and coordinated all bioinformatics activities. O-P.S, L.G.P, and P.A did RNA and DNA library construction and sequencing and participated in genome assembly. O.S did the genome and functional annotation. S.R and P.S participated in genome annotation. O.S analyzed the RNA sequencing data including de novo assembly of RNAseq. O.S did comparative genomics analyses. T.S and N.S were involved in field research for sample isolations. O.S and M.W grown and collected the samples for mass spectrometry. G.L.B, B.B, M.W, and O.S were sequenced were involved in cloning and expression of CYP450 and Lupeol synthase enzymes. N.S and J.L did mass spectrometry, including sample pretreatment, method development, UPLC-HDMS analysis, metabolite identification and data interpretation, and O.S and J.S contributed to data interpretation. O.S and J.S wrote the original manuscript with input from O-P.S , U.R, K.O.

## Acknowledgement

We thank Cory D. Dunn who provided us with yeast strain, and Peter M.J. Burgers, Ville O. Paavilainen, and Juho Kellosalo for giving us the expression vector. We also acknowledge the computational infrastructure of CSC IT Center for Science, Finland. J.S would like to acknowledge the funding from University of Helsinki three-year grant, Academy of Finland (decisions 318288 and 319947), as well as Nanyang Technological University start-up grant.

---

**Supp. Fig 1:**
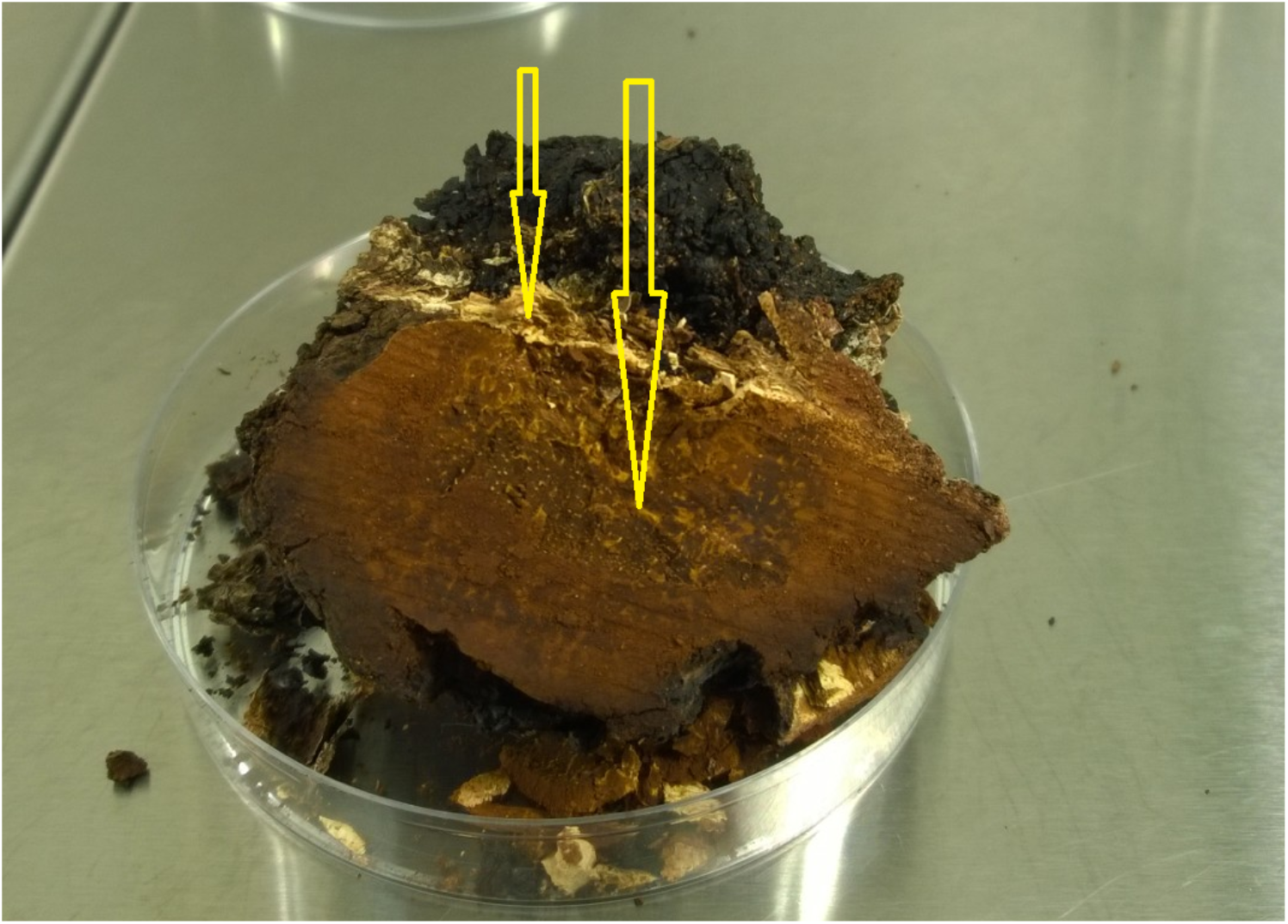
Fragment of *Inonotus obliquus* conk collected from Merikarvia, Finland [N62.00°, E24.74°]. Yellow arrows point at sites where samples were collected for inoculation of culture media. There might be contamination of birch bark at the site where the smaller arrow is pointing. The larger arrow points to the center of the conk where more pure sample was collected.

**Supp. Fig 2:**
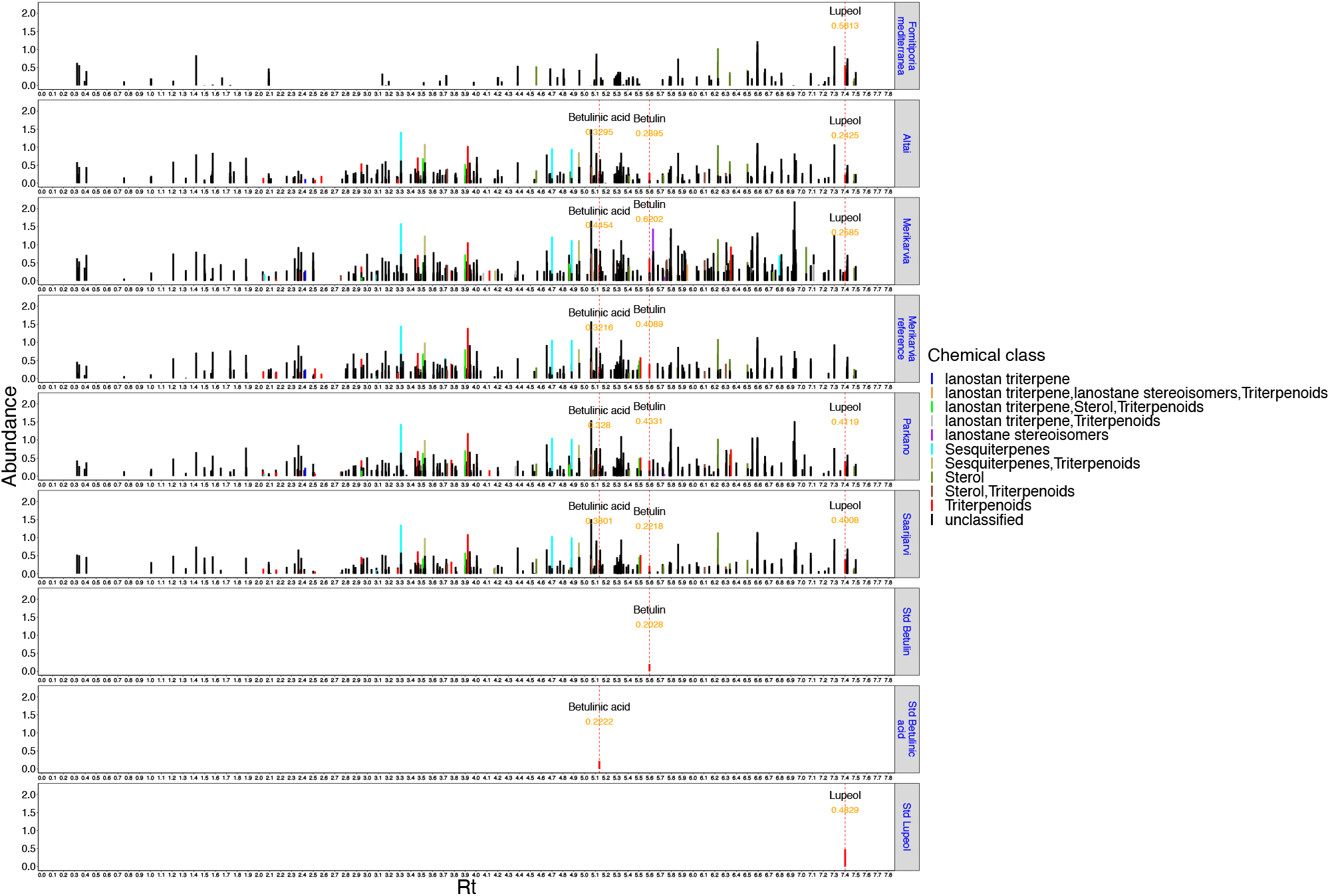
Relative abundances of secondary metabolites among five strains of *I. obliquus*, one strain of *F. mediterranea*, and standard (98%) lupeol, betulin, and betulinic acid. X-axis is the retention time, and Y-axis the relative abundance (ggplot, scales=’free_x’). Vertical red lines show the standard retention times for lupeol, betulin, and betulinic acid, labeled with the name of the compound and its relative abundance. Mass spectra are classified to known chemical class. Std: standard (98%).

**Supp. Fig 3:**
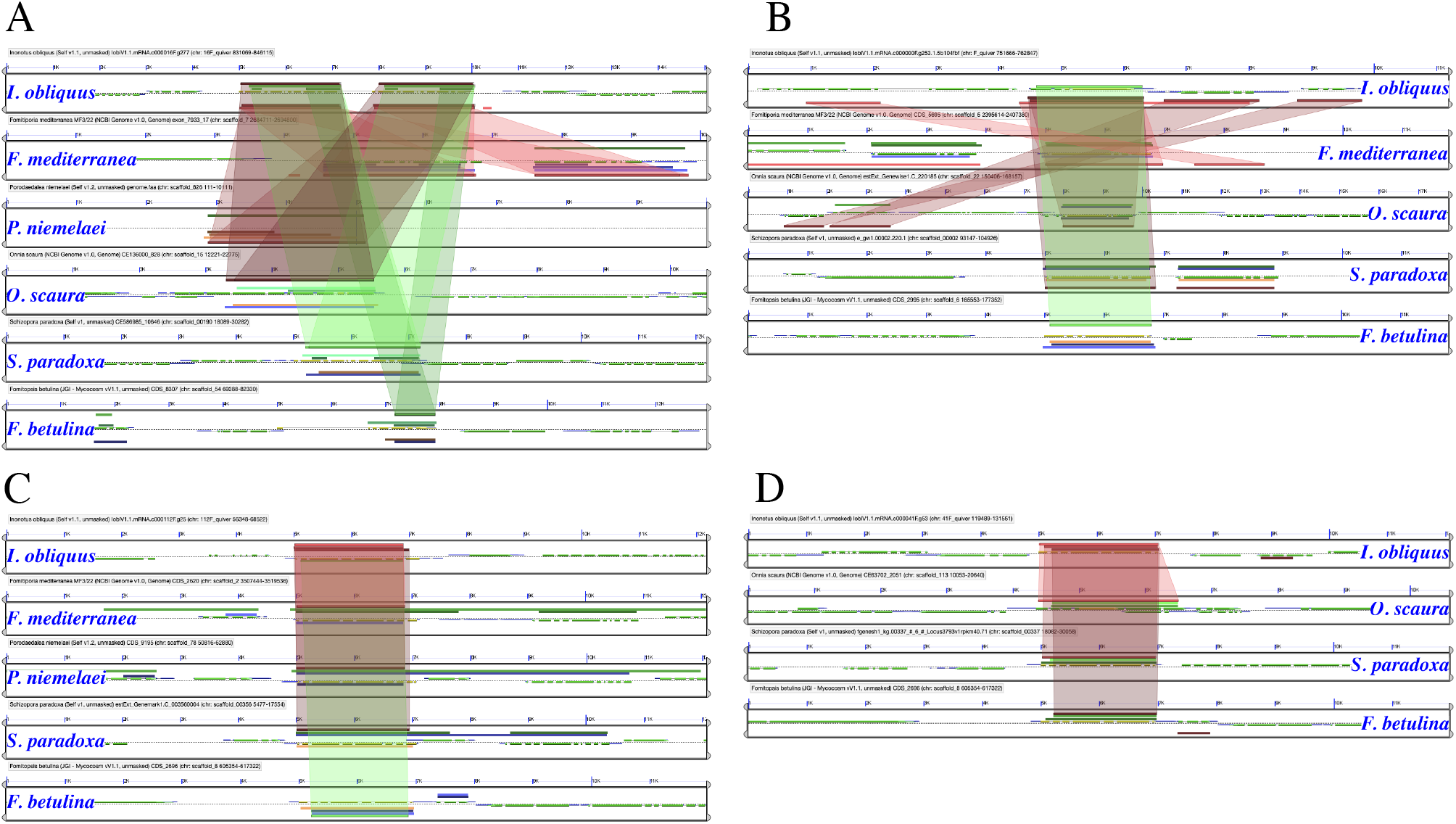
Microsynteny analysis of four cloned genes from *I. obliquus* and four *Hymenochaetales* species and *Fomitopsis betulina*. **A)** gene ID: c000016F.g277, **B)** gene ID: c000000F.g253, **C)** gene ID: c000112F.g25, and **D)** gene ID: c000041F.g53.

**Supp. table 1**: Statistics of I obliquus genome assembly and annotation, genome size of orthologous species, repeat masking statistics, and the GO enrichment of genes adjacent to DNA transposable elements.

**Supp. table 2**: List of putative secreted proteins from I obliquus.

**Supp. table 3**: List of genes from I obliquus which are associated to syntenic or tandemly duplicated regions of the genome.

**Supp. table 4**: List of expanded gene families and their GO enrichment in *I. obliquus*.

**Supp. table 5**: List of homologous C*AZymes from I. obliquus.*

**Supp. table 6**: List of syntenic and tandemly duplicated genes, and gene ontology (GO) enrichment in these genomic regions.

**Supp. table 7**: List of metabolomics fingerprints from five strains of *I. obliquus*, and one strain of *F. mediterranea*.

**Supp. table 8**: List of putative oxygen, heme, ERR-Triad domains across multiple kingdoms.

**Supp. table 9**: List of differentially expressed gene families.

